# In vitro and computational analysis of the putative furin cleavage site (RRARS) in the divergent spike protein of the rodent coronavirus AcCoV-JC34 (sub-genus luchacovirus)

**DOI:** 10.1101/2021.12.16.473025

**Authors:** Annette Choi, Deanndria T. Singleton, Alison E. Stout, Jean K. Millet, Gary R. Whittaker

**Affiliations:** Departments of Microbiology & Immunology, College of Veterinary Medicine, Cornell University, Ithaca, NY, 14853, USA; Public & Ecosystem Health, College of Veterinary Medicine, Cornell University, Ithaca, NY, 14853, USA; Université Paris-Saclay, INRAE, UVSQ, Virologie et Immunologie Moléculaires, Jouyen-Josas, France

## Abstract

The *Coronaviridae* is a highly diverse virus family, with reservoir hosts in a variety of wildlife species that encompass bats, birds and small mammals, including rodents. Within the taxonomic group alphacoronavirus, certain sub-genera (including the luchacoviruses) have phylogenetically distinct spike proteins, which remain essentially uncharacterized. Using in vitro and computational techniques, we analyzed the spike protein of the rodent coronavirus AcCoV-JC34 from the sub-genus luchacovirus, previously identified in *Apodemus chevrieri* (Chevrier’s field mouse). We show that AcCoV-JC34—unlike the other luchacoviruses—has a putative furin cleavage site (FCS) within its spike S1 domain, close to the S1/S2 interface. The pattern of basic amino acids within the AcCoV-JC34 FCS (-RR-R-) is identical to that found in “pre-variant” SARS-CoV-2—which is in itself atypical for an FCS, and suboptimal for furin cleavage. Our analysis shows that, while containing an -RR-R-motif, the AcCoV-JC34 spike “FCS” is not cleaved by furin (unlike for SARS-CoV-2), suggesting the possible presence of a progenitor sequence for viral emergence from a distinct wildlife host.

## Introduction

The animal reservoirs for pandemic potential viruses (including coronaviruses) are focused on the breadth of bat species (order Chiroptera) that exist around the world [1-3]. However certain coronaviruses, notably the sub-genus embecovirus (genus betacoronavirus) currently have no bat-origin examples and have a putative reservoir in animal species within the order Rodentia, which is the most diverse mammalian order on the planet and is well-documented as an important reservoir host for human diseases [4, 5].

While rodents are generally appreciated as an important reservoir for RNA viruses, surveillance and detection of coronaviruses is currently relatively limited. Following the initial discovery of what is now the prototype luchacovirus (Lucheng Rn rat coronavirus, or LRNV), along with two *Betacoronavirus* species [6], a study from Ge *et al*. examined 177 intestinal samples from three species of rodents in Yunnan Province, China and detected both alphacoronaviruses and betacoronviruses in three animal species (*Apodemus chevrieri, Eothenomys fidelis* and *Apodemis ilex*) [7]. Their study reported the full-length genome of a coronavirus (AcCoV-JC34) from *A. chevrieri* (Chevrier’s field mouse) that was designated an alphacoronavirus (sub-genus luchacovirus) based on its genome structure and multiple sequence alignments, which included analysis of the whole genome and the ORF1a/b genes. However, Ge *et al*. noted that both AcCoV-JC34 and LRNV may represent a novel alphacoronavirus species. In particular, they noted that the luchacovirus S gene formed a distinct genetic lineage with low sequence identity (<25%) compared to other well characterized coronaviruses. Ge *at al*. also noted that AcCoV-JC34 S contained two predicted proteolytic cleavage sites, one at residue 508 at the S1/S2 interface, and the other at residue 674 (the fusion peptide-proximal S2’ position).

More recently, a more comprehensive sampling of rodents and other small mammals has identified a diverse range of coronaviruses in such animal reservoirs [8]. To determine the evolutionary history of rodent alphacoronaviruses in more detail, Tsoleridis *et al*. also reported sequence data from viruses sampled from European rodents, to define a single common ancestor for all rodent alphacoronaviruses with a shared recombinant betacoronavirus spike gene—also shared with batCoV HKU2, swine acute diarrhea syndrome (SADS) coronavirus and two shrew coronaviruses [9]. According to Tsoleridis *et al*., the luchacoviruses (including AcCoV-JC34) comprised a distinct lineage within the “recombinant” viruses. In summary, it can be argued that coronaviruses of small mammals, including rodents, are still poorly understood.

We have previously reported that rodent coronavirus AcCoV-JC34 has a weakly predicted furin cleavage site (FCS) is its spike protein [10]. Here, we further analyze the AcCoV-JC34 spike and its “FCS” along with the other luchacoviruses, taking an in vitro and computational perspective.

## Results

### Phylogenetic analysis of luchacoviruses

To understand the relationship of ACoV-JC34 and the other known luchacoviruses, we first constructed a phylogenetic tree of these viruses in comparison to representatives of the diverse coronavirus family, based on spike protein sequences (Figure 1). In agreement with Ge *at al*., luchacoviruses formed a monophyletic group with 100% bootstrap support, indicating a common ancestor origin outside of the established alphacoronavirus branch. Luchacoviruses clustered with rhinacoviruses, which include swine acute diarrhea syndrome coronavirus, Rhinolophus bat coronavirus HKU2, and porcine enteric alpha coronavirus (Figure 1).

**Figure 1.**
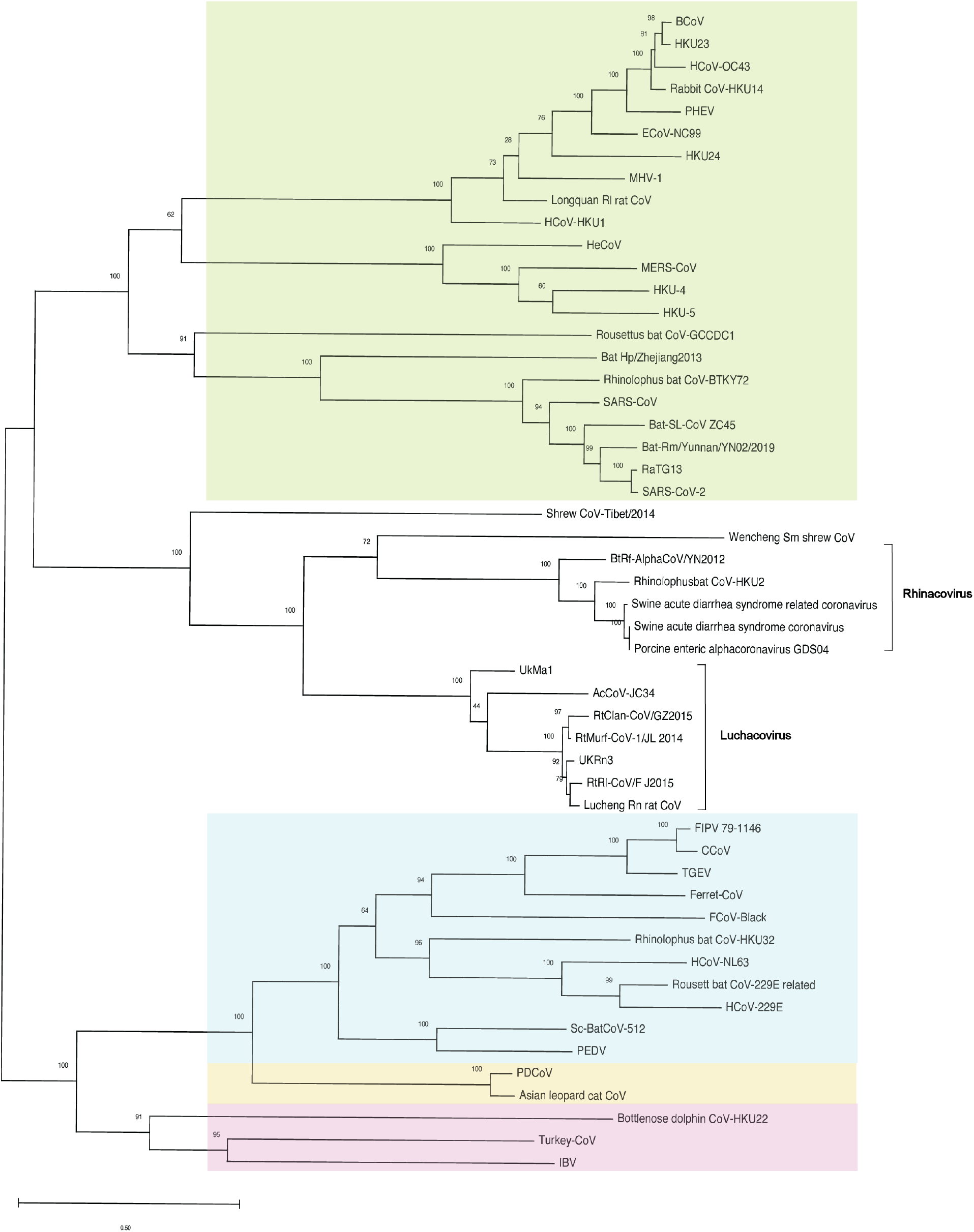
Phylogenetic tree of spike protein sequences. The maximum likelihood phylogenetic tree was constructed using MegaX, 100 bootstraps, from a multiple sequence alignment of the spike sequences. Betacoronavirus spikes are shaded green, alphacoronavirus spikes are shaded blue, deltacoronavirus spikes are shaded yellow and gammacoronavirus spikes are shaded pink

### Geographical distribution of sampled luchacoviruses

The geographical location, dates and rodent species sampled for the currently identified luchacoviruses are summarized in Figure 2 and Table 1. The luchacoviruses sampled to date are from a range of rodent hosts and are from the United Kingdom and several provinces in China (Figure 2), indicating a widespread distribution. Despite being sampled in these distinct locations, as mentioned above, luchacoviruses form a monophyletic group suggesting they have been associated with rodents for an extended period of time.

**Figure 2.**
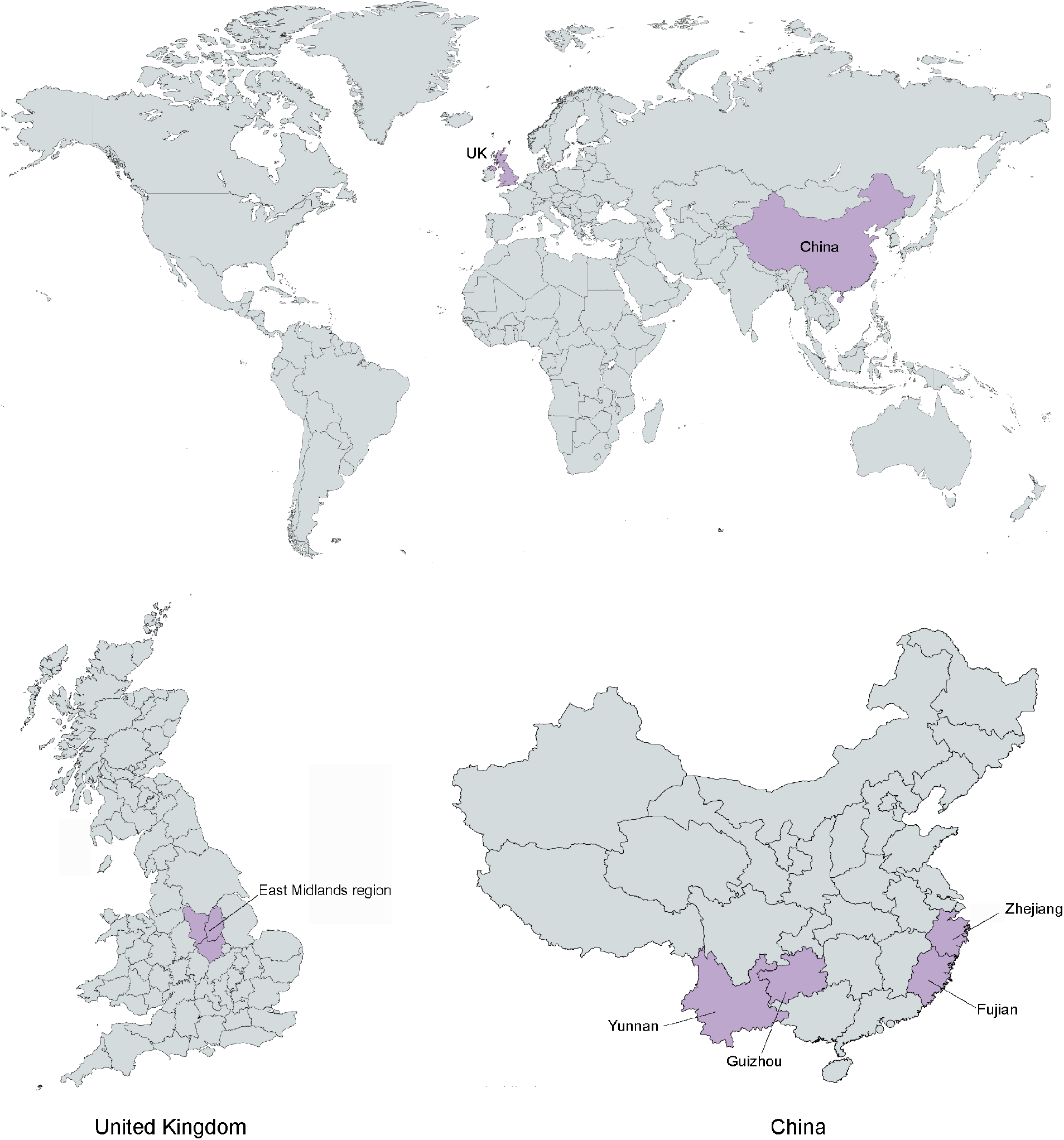
Geographical distribution of currently identified luchacoviruses. Luchacoviruses have been identified from surveillance studies in United Kingdom (East Midlands region) and China (Yuanna, Zhejiang, Fujian, Jilin, and Guizhou provinces).

**Table 1.**
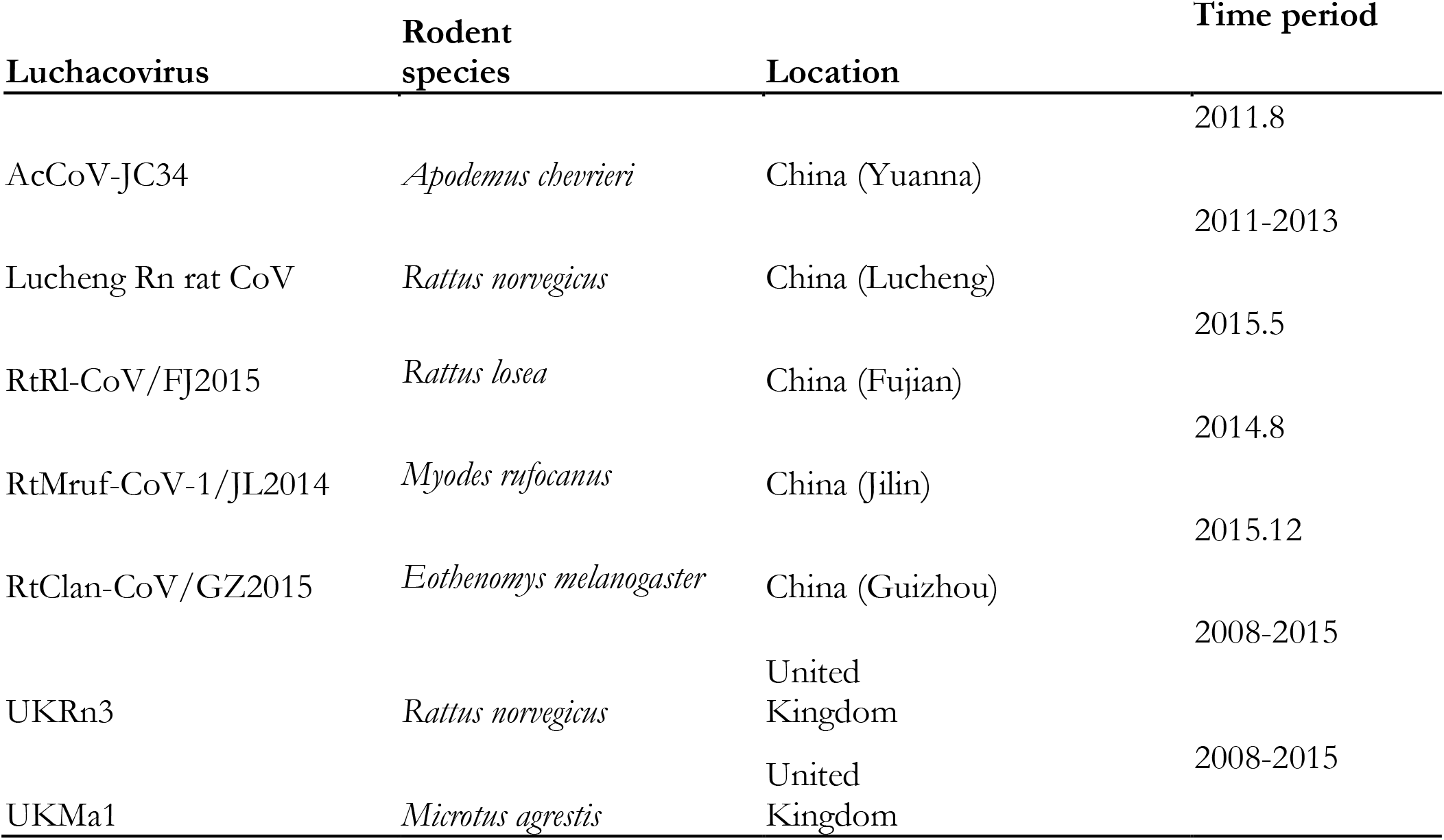
Rodent luchacoviruses identified from surveillance studies.

### Multiple sequence alignment and structural analysis of AcCoV-JC34 spike

A multiple sequence alignment of spike proteins was performed on AcCoV-JC34 spike in comparison to the prototype luchacovirus Lucheng Rn rat CoV (LRNV), as well as SARS-CoV-2, SARS-CoV, HCoV-HKU1, HCoV-OC43 and MERS-CoV. This alignment revealed that the -RR-R-motif present in AcCoV-JC34 does not align precisely with the S1/S2 motif of most coronavirus spikes (Figure 3). However, it aligned with a potential secondary MERS-CoV furin cleavage site (RSTRS).

**Figure 3.**
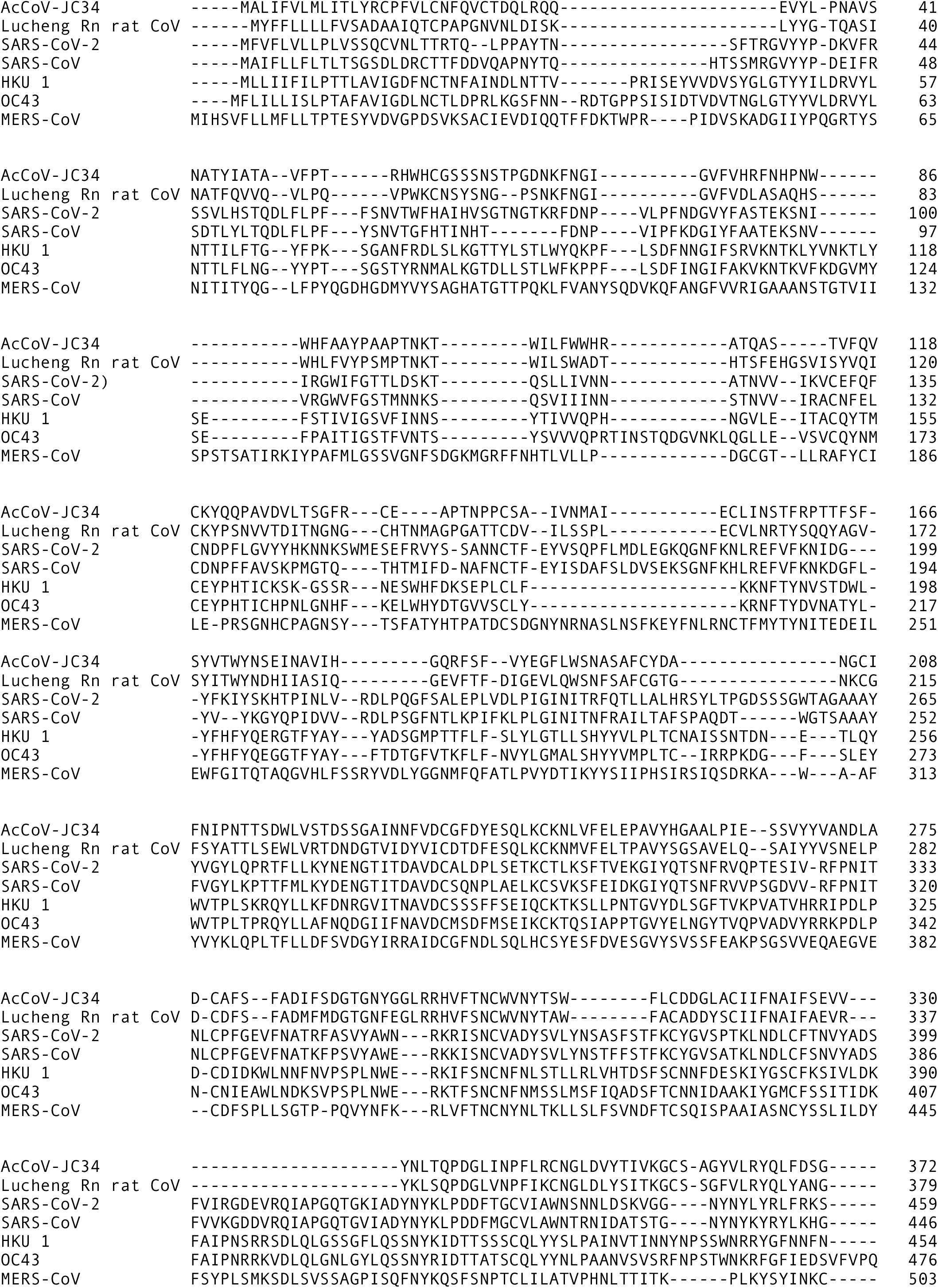
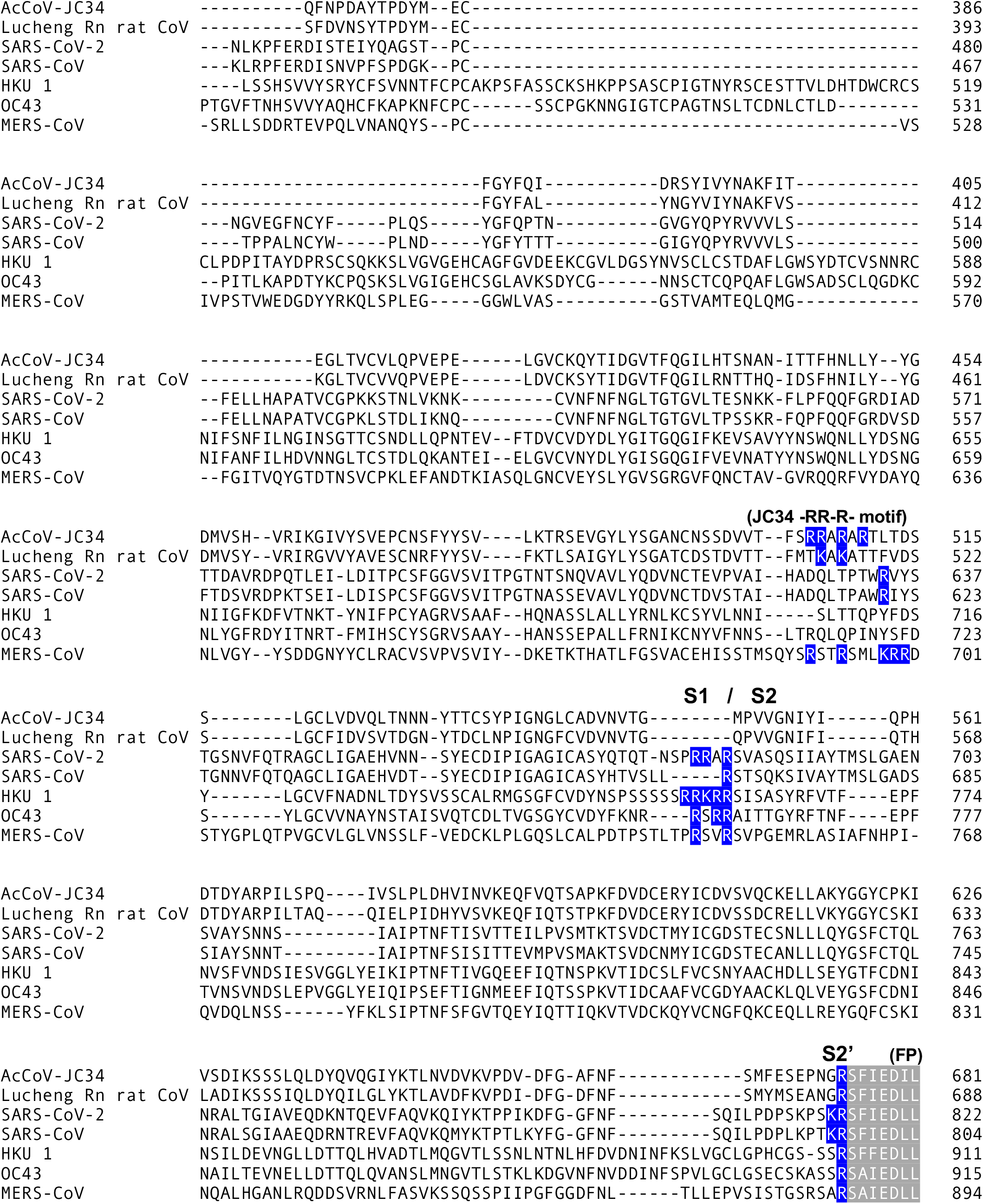
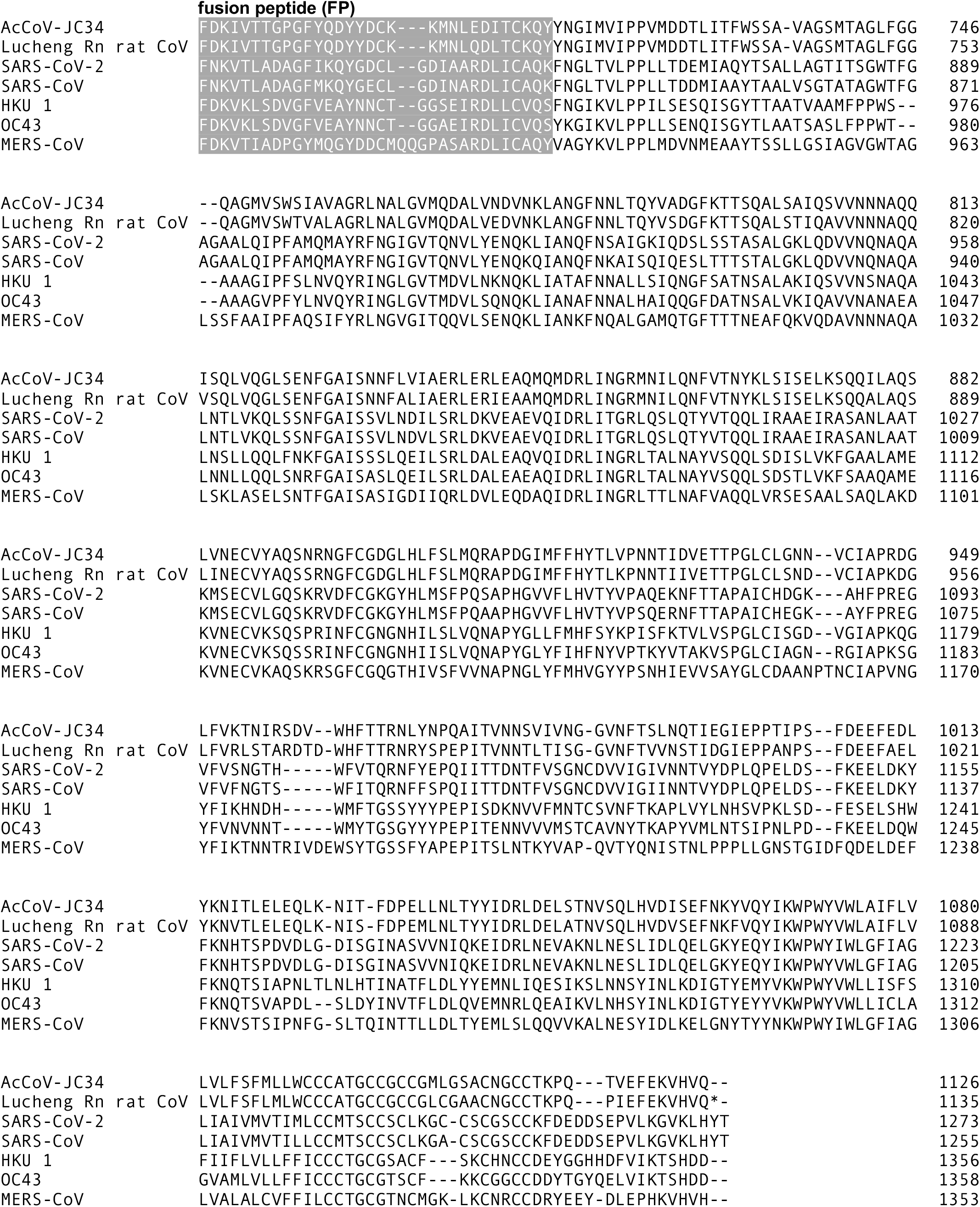

To investigate the structural location of AcCoV-JC34 furin cleavage site, the AcCoV-JC34 spike protein structure was structurally modeled (Figure 4). We used SADS-CoV spike for our modeling due to its available structure in the RCSB protein data bank and relatively high identity with JC34 (41.5%). In our JC34 model, the potential furin cleavage site (-RR-R-) is located in an exposed loop of the protein which is predicted to increases its accessibility to proteases. However, the potential AcCoV-JC34 furin cleavage site was within a loop upstream of the typical S1/S2 furin cleavage site found in other CoVs (see Figure 3). In SARS-CoV-2, this upstream region aligned with a DQLTP sequence upstream of the expected S1/S2 cleavage site.

**Figure 4.**
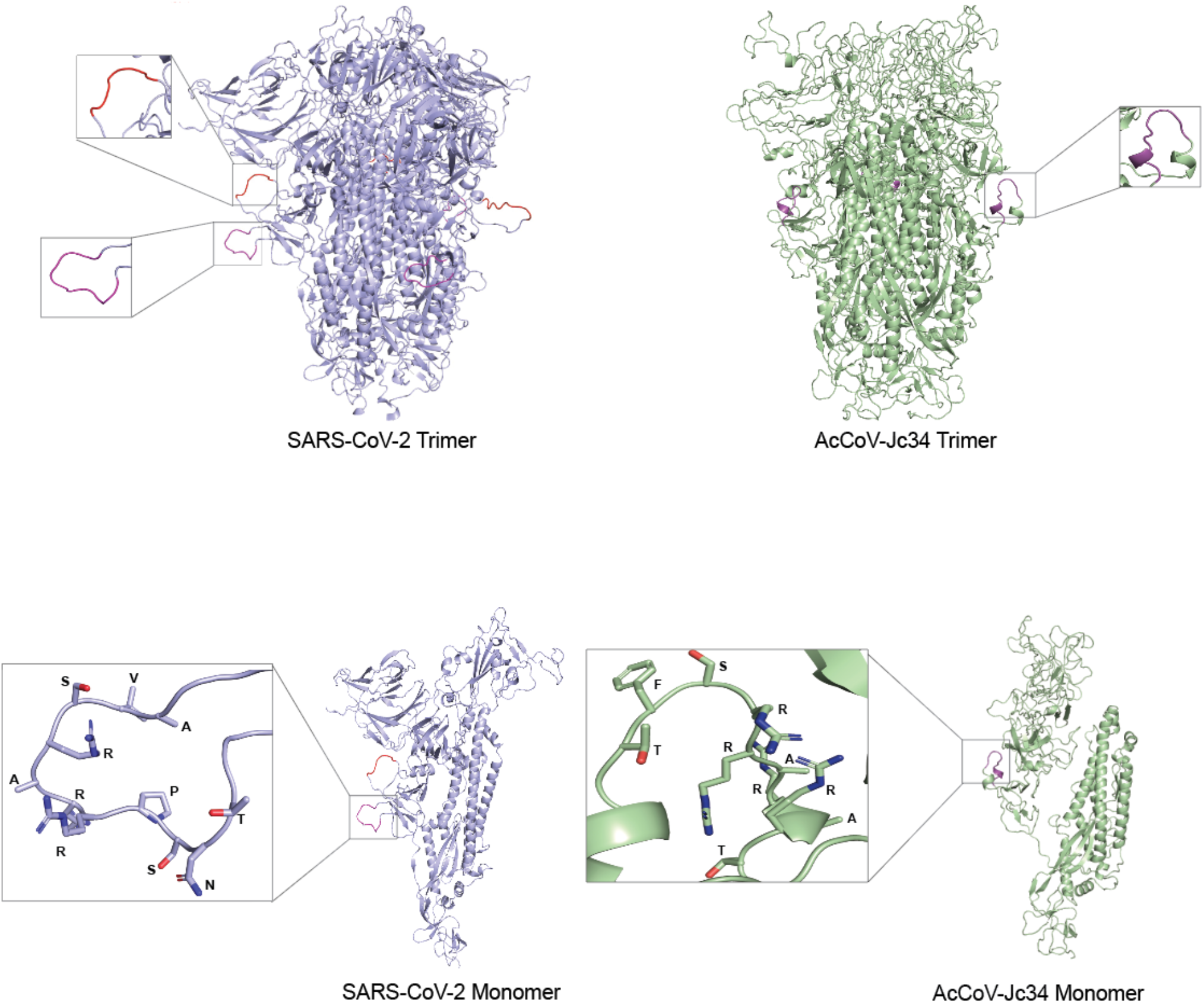
Predicted structure of AcCoV-JC34 spike protein. The AcCoV-JC34 spike protein was modelled based on SADS-CoV spike. In SARS-CoV-2, the pink highlight indicates where the furin cleavage sequence (PRRAR) is located. The red highlight is the location that aligns with AcCoV-JC34 potential furin cleavage site. In the AcCoV-JC34 structural model, the pink highlight indicates location of the potential furin cleavage site (SRRAR).

### Bioinformatic and biochemical analysis of potential AcCoV-JC34 spike cleavage site

To determine whether furin processes the -RR-R-motif in AcCoV-JC34, we first utilized the PiTou and ProP furin cleavage prediction tools (Figure 5). A positive score for Pitou or a score above 0.5 for ProP indicates the likelihood of furin cleavage. AcCoV-JC34 has a weakly predicted furin cleavage site based on the PiTou score (see also [10]). Although bioinformatic tools are useful for prediction, these may not represent biologically relevant cleavage events, which need to be addressed experimentally.

**Figure 5.**
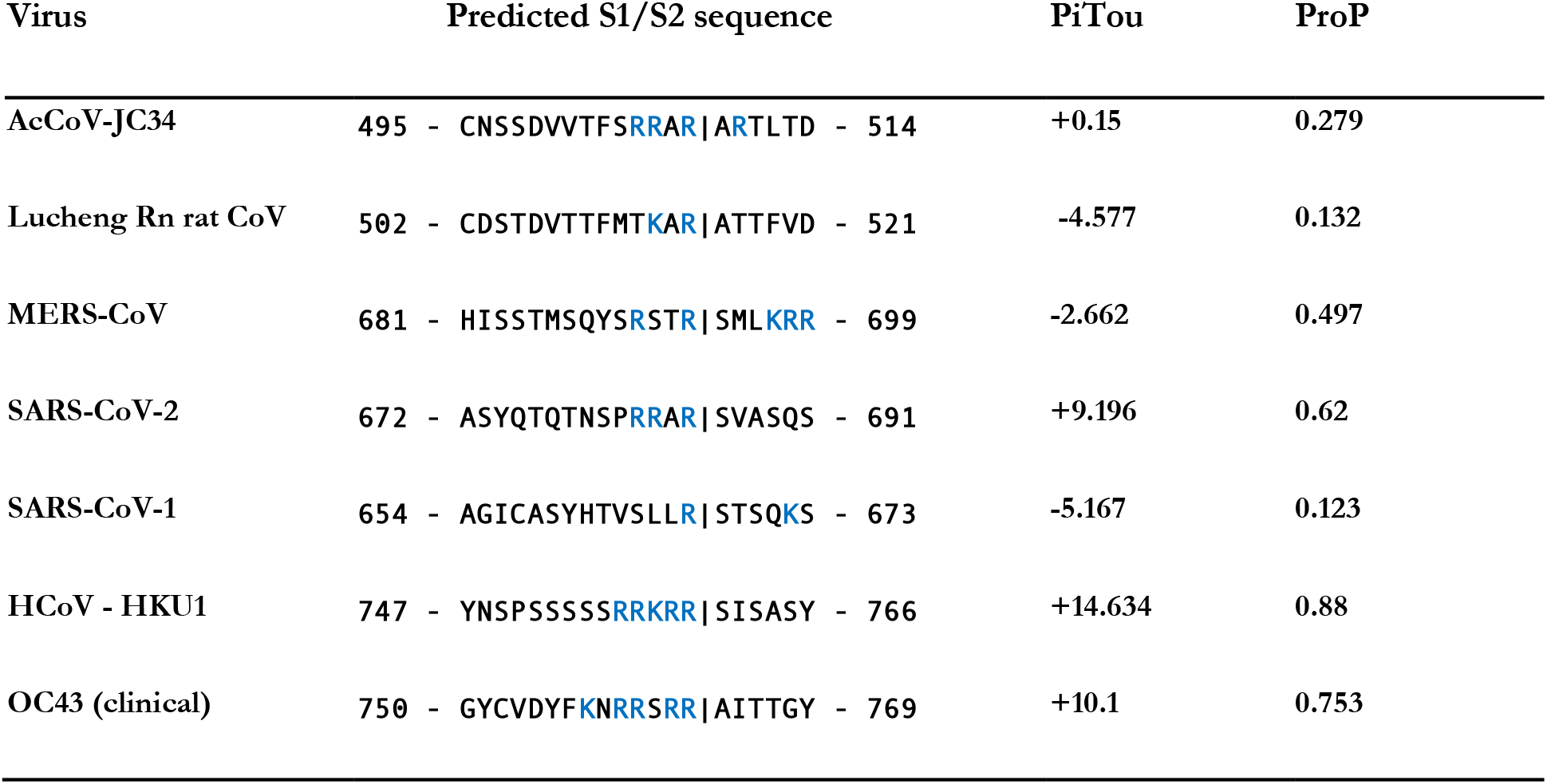
Furin cleavage analysis of CoV S1/S2 cleavage site. CoV S sequences were analyzed using the ProP 1.0 and PiTou 3.0 furin prediction algorithm, generating a score with bold numbers indicating predicted furin cleavage. (|) denotes the position of the predicted S1/S2 cleavage site. Basic resides, arginine (R) and lysine (K), are highlighted in blue.

To directly test whether furin cleaves this site *in vitro*, we performed peptide cleavage assays using furin, along with trypsin as a control. The peptide sequences used were TFMTKARARTTF (Lucheng Rn rat CoV, LRNV), TFSRRARARTL (AcCoV-JC34), and TNSPRRARSVA(SARS-CoV-2). Trypsin cleaved all three peptides with varying efficiency. Furin, as expected from previous studies, cleaved the SARS-CoV-2 peptide; however, it did not cleave the LRNV or JC34 peptides (Figure 6). These data indicate that although AcCoV-JC34 has a minimal furin cleavage sequence (R-X-X-R) it is not able to be cleaved by furin when tested experimentally.

**Figure 6.**
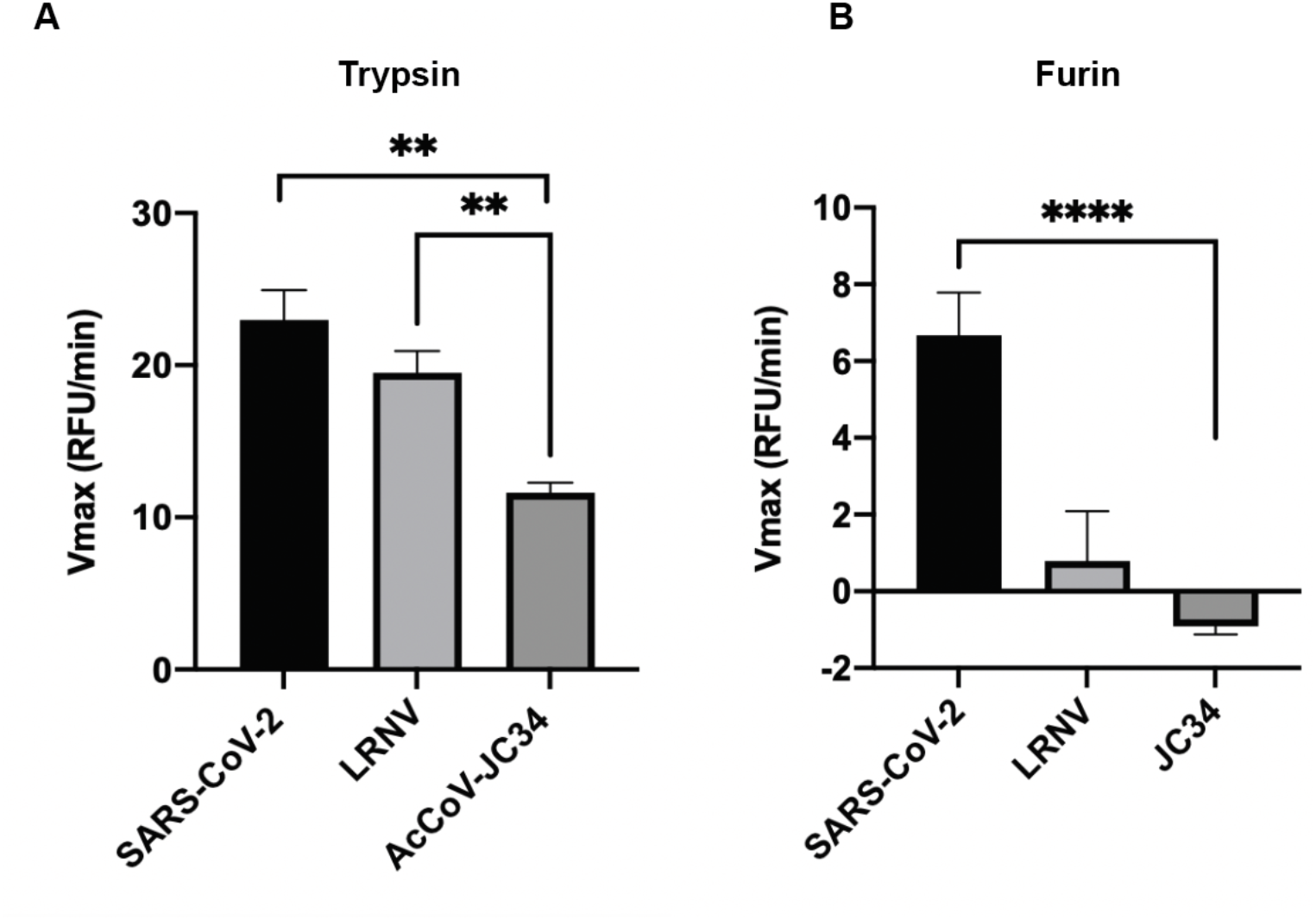
Fluorogenic peptide cleavage assays of the predicted AcCoV-JC34 furin cleavage site. Peptides mimicking the S1/S2 site of the SARS-CoV-2 WT, AcCoV-JC34, and Lucheng Rn rat CoV (LRNV) were evaluated for in vitro cleavage with A) trypsin and B) furin proteases under pH 7.4 (trypsin), and 7.5 (furin) conditions. Trypsin cleaved all three peptides, while furin only cleaved SARS-CoV-2.

## Discussion

The “furin cleavage site” or FCS of SARS-CoV-2 has been at the center of the many discussions on the origin of the COVID-19 pandemic; see [11] for a recent summary. Despite being interpreted as “highly unusual”, an FCS is—to the contrary—very common among the *Coronaviridae* [12], with sarbecoviruses and most alphacoronaviruses being the exception rather than the rule in lacking this important regulatory sequence. In fact, many zoonotic coronaviruses and those in reservoir hosts appear to contain sequences and structural loops at the S1/S2 interface that are sub-threshold for furin-mediated cleavage [13-16] and may be “poised” for spillover events. Examples include “pre-variant” SARS-CoV-2, as well as the sarbecoviruses RmYN02, RacCS203, BANAL-20-116, BANAL-20-246 that have potential phylogenetic homology to the SARS-CoV-2 FCS [17]—and may include the luchacovirus AcCoV-JC34 analyzed here. It is noteworthy that AcCoV-JC34 is the only luchacovirus containing this -R-RR-motif.

While containing an -RR-R-motif, as found in SARS-CoV-2, the data presented here show that this AcCoV-JC34 sequence is not cleaved by furin. The reasons for this are currently unclear. One possibility is that the upstream proline found in SARS-CoV-2, as well as in other spike cleavage site sequences, may promote cleavage by creating a structural turn beneficial for furin activity. It is also possible that the additional downstream arginine residue in AcCoV-JC34 spike may be inhibitory for the tight active site binding pocket present in furin [18]. Alternatively, the structural loop present in AcCoV-JC34 spike may be cleaved by other proprotein convertases of the furin family that have less stringent cleavage requirements, or by trypsin-like enzymes or cathepsins. Notably, the -RR-R-motif is rare in furin substrates, and only other known example of this sequence motif in FurinDB (a database of furin substrates) is found in proaerolysin, a bacterial toxin [19].

One notable aspect of the -RR-R-motif in AcCoV-JC34 is that is does not align precisely with the S1/S2 motif of most coronavirus spikes (see Figure 3) and is a structurally exposed location above the typical S1/S2 loop (see Figure 4). Analysis of the MERS-CoV spike also shows an addition putative FCS in the MERS-CoV spike (SRSTRS); while this contains a minimal furin motif this sequence shows low scores for furin cleavage with both Pitou and ProP, and FRET-based peptides were not cleaved by furin in biochemical cleavage assays—in contrast to the PRSVRS motif at the expected S1/S2 junction (J. K. Millet, unpublished results). Nevertheless, it is possible that, as with AcCoV-Jc34, this “secondary” MERS-CoV sequence comprises a “blocked” FCS due to flanking hydrophobic and charges residues in the downstream C-terminal positions (i.e., SRSTRSMLKRRDS). This putative secondary cleavage site also lacks an upstream proline/proline-rich region, as with many other S1/S2 regions that are known to be cleaved by furin.

For SARS-CoV-2, it is clear that selection is occurring to up-regulate the spike FCS, as seen with several of the highly transmissible variants that have emerged [20-24]. The FCS can also be readily down regulated upon Vero cell adaptation; for examples see refs [25, 26]. Likewise, some coronaviruses in animal reservoirs may be “poised” for proteolytic cleavage-activation at S1/S2, with selection occurring along with modifications to their receptor binding domain. One interesting example of this may exemplified by the MERS-like bat-CoVs HKU-4 and HKU-5, with HKU-4 binding human DPP4, but having no identifiable FCS, and with HKU-5 not able to bind hDPP4 and having a robust FCS [27].

Our studies highlight the possible presence of a distinct proteolytic cleavage loop in the coronavirus spike protein and the specific features of the luchacovirus spike—which along with that found in the rhinacoviruses (e.g., SADS-CoV) appears to represent an evolutionary disparate spike protein with apparent similarities to a betacoronavirus spike protein (see Figure 1), despite the taxonomic designation of these viruses as alphacoronaviruses.

## Methods

### Furin prediction calculations

Prop: CoV sequences were analyzed using the ProP 1.0 Server hosted at: cbs.dtu.dk/services/ProP/. PiTou: CoV sequences were analyzed using the PiTou V3 software hosted at: http://www.nuolan.net/reference.html.

### Amino acid alignments and phylogenetic trees

Multiple sequence alignment was performed on coronavirus spike protein using Geneious Prime ® (v.2019.2.3. Biomatters Ltd.). A maximum likelihood phylogenetic tree was constructed using MegaX(, 100 boot strap replicates based on the spike protein. Amino acid sequences of S were obtained from NCBI GenBank. Accession numbers are: AcCoV-Jc34 (YP_009380521), Asian leopard cat CoV (EF584908.1),Bat_Hp/Zhejiang2013(YP_009072440),Bat-Rm/Yunnan/YN02/2019 (QPD89843.1), Bat-SL-CoV_ZC45 (AVP78031.1), BCoV (P15777), Bottlenose dolphin CoV-HKU22 (AHB63508), BtRf-AlphaCoV/YN2012 (YP_009200735), CCoV (AY436637.1), ECoV-NC99 (AAQ67205.1), FCoV-Black (EU186072.1), Ferret-CoV (NC_030292.1), FIPV 79-1146 (DQ010921.1),HCoV-229E(NC_002645.1),HCoV-HKU1(NC_006577),HCoV-NL63(NC_005831.2),HCoVOC43(NC_006213.1),HeCoV(MK679660.1),HKU4(YP_001039953),H KU5(YP_001039962), HKU23(QEY10673),HKU24(QOE77327), IBV(NC_001451.1), Longquan Rl rat CoV (QOE77336.1), Lucheng Rn rat CoV(QOE77268.1), MERS-CoV(AFS88936.1), MHV-1 (ACN89742), PDCoV (MN942260.1), PEDV(NC_003436.1), PHEV(QTF73995.1), Porcine enteric alphacoronavirus GDS04 (ASK51717.1), Rabbit CoV-HKU14(AFE48827), RaTG13(QHR63300), Rhinolophus bat coronavirus HKU2 (YP_001552236.1), Rhinolophus bat CoV-BTKY72 (APO40579.1), Rhinolophus bat CoV-HKU32 (QCX35178), Rhinolophusbat CoV-HKU2 (YP_001552236.1), Rousett bat CoV-229E related(QHA24665), Rousettus bat CoV-GCCDC1(QKF94914), RtClan-CoV/GZ2015 (), RtMurf-CoV-1/JL 2014 (ATP66738), RtRl-CoV/F J2015 (KY370050), SARS-CoV (AAT74874.1), SARS-CoV-2 Wuhan-Hu-1 (YP_009724390.1), Sc-BatCoV-512 (ABG47078), Swine acute diarrhea syndrome coronavirus(AVM41569.1), Swine acute diarrhea syndrome related coronavirus (AVM80500.1), TGEV(P07946), Turkey-CoV (QRR19172), UkMa1(QBG64648), UKRn3(QBG64657).

### Spike structural modelling

Pairwise amino acid alignment between AcCoV-Jc34 (YP_009380521) and SADS-CoV (AVM80500) was performed using Geneious Prime ® (v.2019.2.3. Biomatters Ltd.). S protein models were built based on the SADS-CoV structure obtained from RCSB (PDB: 6M39), using UCSF Chimera (v.1.14, University of California) through the modeler homology tool of the Modeller extension (v.9.23, University of California).

### Fluorogenic peptide cleavage assays

Fluorogenic peptide cleavage assays were performed as described previously [14]. Each reaction was performed in a 100 μL volume consisting of buffer, protease, and AcCoV-Jc34 (TFSRRARARTL) or Lucheng Rn rat CoV (TFMTKARARTTF) or SARS-CoV-2 S1/S2 WT (TNSPRRARSVA) fluorogenic peptide in an opaque 96-well plate. For trypsin catalyzed reactions, 0.8 nM/well TPCK trypsin was diluted in PBS buffer. For furin catalyzed reactions, 1 U/well recombinant furin was diluted in buffer consisting of 20 mM HEPES, 0.2 mM CaCl2, and 0.2 mM β-mercaptoethanol, at pH 7.0. Fluorescence emission was measured once per minute for 60 minutes using a SpectraMax fluorometer (Molecular Devices) at 30 °C with an excitation wavelength of 330 nm and an emission wavelength of 390 nm. Vmax was calculated by fitting the linear rise in fluorescence to the equation of a line.

## Acknowlegements

DTS is supported by the Howard Hughes Medical Institute-Cornell University Transfer (HHMI-CURT) program

AES was supported by NIH Comparative Medicine Training Program T32OD011000.

Work in the author’s lab is funded in part by the National Institute of Health research grant R01AI35270 (to GW).

We thank the members of the Whittaker Lab, past and present, for their helpful discussions during the preparation of this manuscript.

